# Mechanosensitive calcium signaling in filopodia

**DOI:** 10.1101/2020.10.21.346247

**Authors:** Artem K. Efremov, Mingxi Yao, Michael P. Sheetz, Alexander D. Bershadsky, Boris Martinac, Jie Yan

**Author notes:** These authors equally contributed to the study (Artem K. Efremov, Mingxi Yao). Correspondence and requests for materials should be addressed to: A.K.E. and J.Y.

## Abstract

Filopodia are ubiquitous membrane projections that play crucial role in guiding cell migration on rigid substrates and through extracellular matrix by utilizing yet unknown mechanosensing molecular pathways. As recent studies show that Ca^2+^ channels localized to filopodia play an important role in regulation of their formation and since some Ca^2+^ channels are known to possess mechanosensing properties, activity of filopodial Ca^2+^ channels might be tightly interlinked with the filopodia mechanosensing function. We tested this hypothesis by monitoring changes in the intra-filopodial Ca^2+^ level in response to application of stretching force to individual filopodia of several cell types. It has been found that stretching forces of tens of pN strongly promote Ca^2+^ influx into filopodia, causing persistent Ca^2+^ oscillations that last for minutes even after the force is released. Most of the known mechanosensitive Ca^2+^ channels, such as Piezo 1, Piezo 2 and TRPV4, were found to be dispensable for the observed force-dependent Ca^2+^ influx. In contrast, L-type Ca^2+^ channels appear to be a key component in the discovered phenomenon. Since previous studies have shown that intra-filopodial transient Ca^2+^ signals play an important role in guidance of cell migration, our results suggest that the force-dependent activation of L-type Ca^2+^ channels may contribute to this process. Overall, our study reveals an intricate interplay between mechanical forces and Ca^2+^ signaling in filopodia, providing novel mechanistic insights for the force-dependent filopodia functions in guidance of cell migration.

**Significance statement:** We found that tensile forces of tens of pN applied to individual filopodia trigger Ca^2+^ influx through L-type Ca^2+^ channels, producing persistent Ca^2+^ oscillations inside mechanically stretched filopodia. Resulting elevation of the intra-filopodial Ca^2+^ level in turn leads to downstream activation of calpain protease, which is known to play a crucial role in regulation of the cell adhesion dynamics. Thus, our work suggests that L-type channel-dependent Ca^2+^ signaling and the mechanosensing function of filopodia are coupled to each other, synergistically governing cell adhesion and motion in a force-dependent manner. Since L-type Ca^2+^ channels have been previously found in many different cell types, such as neural or cancer cells, the above mechanism is likely to be widespread among various cell lines.

## Introduction

The process of cell migration plays the central role in development and maintenance of multicellular organisms. Wounds healing, immune response to exogenous pathogens, embryonic tissue morphogenesis – these are just a few examples of a large number of vital biological processes that rely on highly ordered collective cell migration, which is required for proper functioning of living organisms (1–4). To guide their motion through extracellular matrix (ECM), living cells use dynamic membrane projections known as filopodia, which are responsible for mechanical and chemical sensing of the surrounding microenvironment as well as formation of initial adhesion contacts with ECM or other cells (1–5). For example, it has been recently shown that filopodia play the central role in guiding cell durotaxis, a preferential cell movement along the gradient of the substrate rigidity (6, 7). Furthermore, numerous experimental studies suggest that abnormally high filopodia activity is a typical feature of aggressive cancer cells that results in their high motility, leading to metastases (3). Thus, understanding molecular mechanisms that regulate filopodia dynamics and adhesion may provide important insights into the cell migration process and its alteration during neoplastic development.

Recently, several proteins necessary for filopodia formation, growth and adhesion have been identified. Filopodia are actin-rich membrane extensions of ~ 2-40 μm in length that were shown to contain a bundle (~ 12-20) of parallel actin filaments crosslinked with each other by fascin proteins (2, 3, 5, 8–10). Polymerization of these filaments is carried out by synergistic cooperation between formins [mDia1,2 (11, 12) and FMNL2,3 (13–15)] and actin uncapping Ena/VASP proteins (16–18), which localize to the dynamic ends of the actin filaments at the filopodia tips. Filopodia adhesion to ECM is mediated by integrin molecules, which can be linked to actin filaments via talin proteins (19, 20). Talins in addition have binding sites for RIAM proteins that provide connection to VASP/profilin complexes, promoting polymerization of actin filaments (20–22). Furthermore, filopodia also contain actin-based molecular motors, such as myosin X, which facilitates filopodia formation, and promotes activation and/or transportation of integrin and VASP proteins (23–27), see schematic Figure 1A showing the main filopodial components.

**Figure 1.**
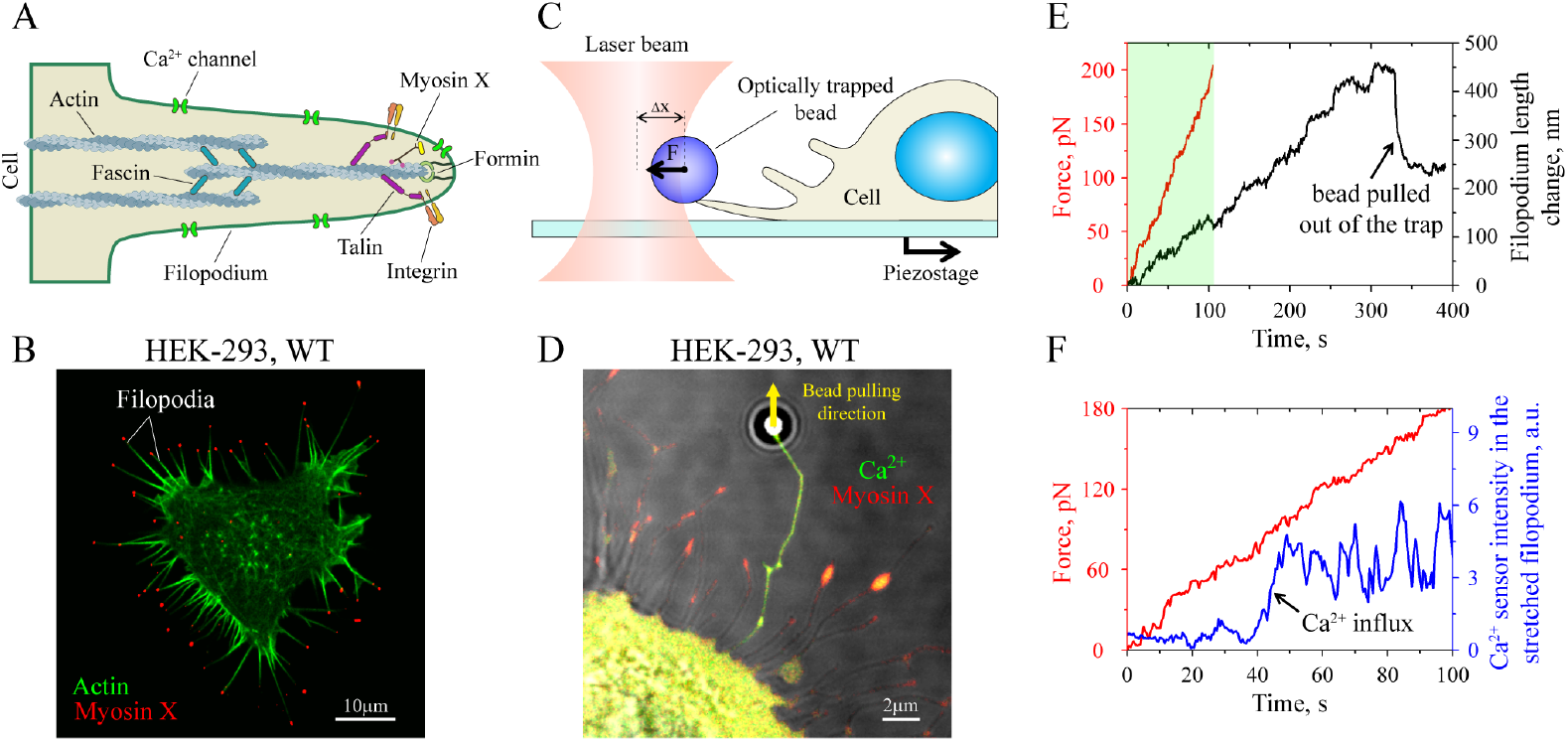
Filopodia components and experimental setup for filopodia stretching. **A.** Schematic illustration of a filopodium and its several known key components. **B.** Typical view of a wild-type (WT) HEK-293 cell transfected with mApple-myosin X construct, which induces filopodia formation. In the figure, mApple-myosin X, which usually clusters at the filopodia tips, is shown in red color; whereas, actin filaments, which are labeled with F-tractin-GFP, are shown in green. **C.** Schematic illustration of the optical tweezers’ experimental setup. Optically trapped microbeads coated with either fibronectin or concanavalin A (ConA) were used to form an adhesion contact with the tip of a filopodium. Filopodium stretching was commenced by moving the microscope stage with the cell body away from the axis of the laser beam. This resulted in generation of a pulling force, *F,* on the filopodium tip. **D.** Fibronectin-coated microbead attached to the filopodium tip pulled by the optical trap away from the cell edge. Filopodium stretching-induced Ca^2+^ signal indicated by GCaMP6f Ca^2+^ sensor (shown in green) as well as mApple-myosin X (shown in red) can be clearly seen from the figure. **E.** Representative time courses of the pulling force and the length change of a stretched filopodium when the cell is moved away from the optical trap at a speed of ~ 5 nm/s. **F.** Time-dependent changes in the pulling force applied to the filopodium (data in red) and the Ca^2+^ sensor intensity (data in blue) corresponding to the experiment shown in panel E.

Although the major proteins contributing to the filopodia formation, growth and adhesion have been identified, there is still a large gap in understanding of how their emergent collective behaviour results in the filopodia’s ability to guide cell migration. Specifically, recent studies show that filopodia can probe and sense mechanical properties of the surrounding environment, guiding cell migration towards stiffer substrate (6, 28). Consistently, a recent work reported that a few pN forces applied to the tips of filopodia can significantly promote the filopodia adhesion and growth in HeLa-JW cells (7). Yet, the exact molecular mechanisms that underlie filopodia’s mechanosensitivity still remain unclear.

Interestingly, it has been recently shown that Ca^2+^ channels are also required for proper formation and stabilization of filopodia in many different cell types (29). Furthermore, it was found that filopodia-dependent transient Ca^2+^ signals play a major role in guidance of cell migration (30). Importantly, several membrane ion channels have been also previously suggested to contribute to cells’ mechanosensing either by direct response to the membrane tension or to tensile forces originating from the cell cytoskeleton. Of particular interest are Ca^2+^-conducting channels such as Piezo 1 and Piezo 2 as well as members of TRP family (31, 32). Upon application of mechanical perturbations to a cell, these channels become rapidly activated through direct or indirect molecular mechanisms, causing influx of extracellular Ca^2+^ into the cell. The latter triggers activation of diverse downstream signaling cascades, many of which play critical roles in such important physiological processes as touch sensation, stem cell differentiation, development of vasculature and various human-related diseases (31, 32). Together with previous works that revealed filopodia as mechanosensing cell structures, these studies point to a possibility that transient Ca^2+^ signalling and the mechanosensing function of filopodia are coupled to each other, synergistically governing cell motion in a force-dependent manner.

Based on the above reasoning, we hypothesized that transient Ca^2+^ signals generated by filopodia may depend on mechanical stretching of the latter. To test this hypothesis, we used previously reported optical tweezers setup (7) to stretch individual filopodia over a physiological range of forces and examined whether the filopodia-related Ca^2+^ signals experience any change in response to the applied mechanical load. It has been found that filopodia produce persistent Ca^2+^ oscillations in a force-dependent manner, which on a large part result from Ca^2+^ influx through voltage-gated L-type Ca^2+^ channels that have been previously discovered in many cell types, including muscle, glial, neuronal and cancer cells (29, 33). Furthermore, by using a calpain activation sensor, it has been shown in our study that the force-induced influx of Ca^2+^ into mechanically stretched filopodia results in downstream activation of calpain protease, which is known to be involved in regulation of the cell adhesion dynamics (34). Altogether, our results suggest that L-type Ca^2+^ channel-dependent signaling plays an important role in mechanosensing function of filopodia, regulating cell adhesion dynamics and motion in a force-dependent manner.

## Results

### Mechanical stretching of individual filopodia activates intra-filopodial Ca^2+^ oscillations

First, we studied how mechanical forces alter Ca^2+^ signaling of filopodia in several cell lines. To promote filopodia formation in those cells, we used either mApple-myosin X construct or constitutively-active GFP-Cdc42 (Q61L), which are the two filopodia growth regulators known to be frequently overexpressed in multiple human cancers (35, 36). Figure 1B shows a representative image of a wild-type (WT) HEK-293 cell transfected with mApple-myosin X. Numerous filopodia of several micrometers (μm) length with myosin X-enriched tips can be clearly seen from the image.

In order to visualize changes in the intra-filopodial Ca^2+^ level, we co-transfected HEK-293 WT cells with mApple-myosin X construct and GCaMP6f Ca^2+^ sensor (37). Interestingly, it has been found that filopodia of such cells occasionally produce Ca^2+^ bursts (see Movie 1). However, frequency of these events appeared to be very low – measurements showed that only ~ 12 % of filopodia (i.e., 11 filopodia out of N = 92 monitored) generated 1 or 2 short bursts of transient Ca^2+^ signals lasting only for ~ 18 ± 5 s (mean ± s.e.m.) during 5-6 min observation period.

To check whether attachment of fibronectin-coated polystyrene microbeads to filopodia has any effect on Ca^2+^ signal behavior, we used optical tweezers to put microbeads onto the tips of individual filopodia, holding them there for ~ 2-3 sec to initiate beads interaction with filopodia before turning the trap off (see schematic Figure 1C). Such simple binding of microbeads to filopodia alone in the absence of applied mechanical load did not change significantly the filopodial Ca^2+^ firing rate – only 25% of filopodia with microbeads produced a single transient Ca^2+^ signal during ~ 5-6 min observation period (the total number of monitored beads was N = 8), see Movie 1.

Interestingly, all the beads attached to filopodia moved in the direction towards the cell body at a rate of 26.9 ± 2.5 nm/s (mean ± s.e.m.), which is similar to the rate of centripetal movement of actin filaments inside filopodia (7, 38). This indicated that fibronectin-coated microbeads were likely engaged to the filopodia actin cytoskeleton, which caused the beads movement towards the cell body due to the retrograde actin flow. Similar behavior was also observed in the case of concanavalin A-coated microbeads (ConA), suggesting that such movement of microbeads towards the cell body may not necessarily depend on the specific beads’ interaction with integrin complexes.

Next, we investigated the role of extracellular mechanical forces in filopodia-dependent generation of Ca^2+^ signals. Optical tweezers setup schematically shown in Figure 1C was used to apply 0-200 pN forces to fibronectin-coated microbeads adhered to the filopodia tips (see Methods section for more details). The pulling force was generated by movement of the microscope piezo-stage away from the optical trap axis at a constant speed of ~ 5 nm/s, stretching the filopodium attached to the trapped microbead with gradually increasing force.

In the presence of mechanical stretching, intense and highly dynamic Ca^2+^ signals were observed in ~ 82% of pulled filopodia (9 out of N = 11 tested filopodia from different cells) that lasted for many minutes, which was in strong contrast to rare Ca^2+^ signals produced by unperturbed filopodia (~ 12%, 7 out of N = 57 filopodia) or by filopodia bound to microbeads in the absence of applied mechanical load (~ 25%, 2 out of N = 8 filopodia). Figure 1D and Movies 2A and 2B show a typical example of a mechanically stretched filopodium with a clearly visible force-induced Ca^2+^ signal. Interestingly, from Movie 2B it can be seen that the Ca^2+^ signal was propagating from the tip of the stretched filopodium towards the cell body, which was a general trend in the pulled filopodia.

Figures 1E-F display representative time traces of the force applied to a stretched filopodium, and the corresponding changes in the filopodium length and intra-filopodial Ca^2+^ level. As can be seen from Figure 1F, the Ca^2+^ sensor intensity increased abruptly at ~ 45 sec after the beginning of the filopodium pulling when the force reached ~ 70 pN level. Measurements from eight independent experiments indicate that the average force required for activation of the Ca^2+^ signal inside filopodia is *F*activation = 78 ± 22 pN (mean ± s.e.m), and it does not show any apparent correlation with the extension change of the pulled filopodia (Figure S1).

Once the Ca^2+^ signal appeared, it could persist for several minutes regardless of whether the force was retained or released (see Figure 2C, middle panel), suggesting that filopodia-dependent Ca^2+^ signaling system has a memory effect, and mechanical stretching is required only for initiation of the first several Ca^2+^ impulses. Furthermore, in ~ 73% of the cases (8 out of N = 11 tested filopodia), during the period when the Ca^2+^ signal persisted, the strength of the signal oscillated in time with a period of ~ 10 s (Figure 2C, right panel).

**Figure 2.**
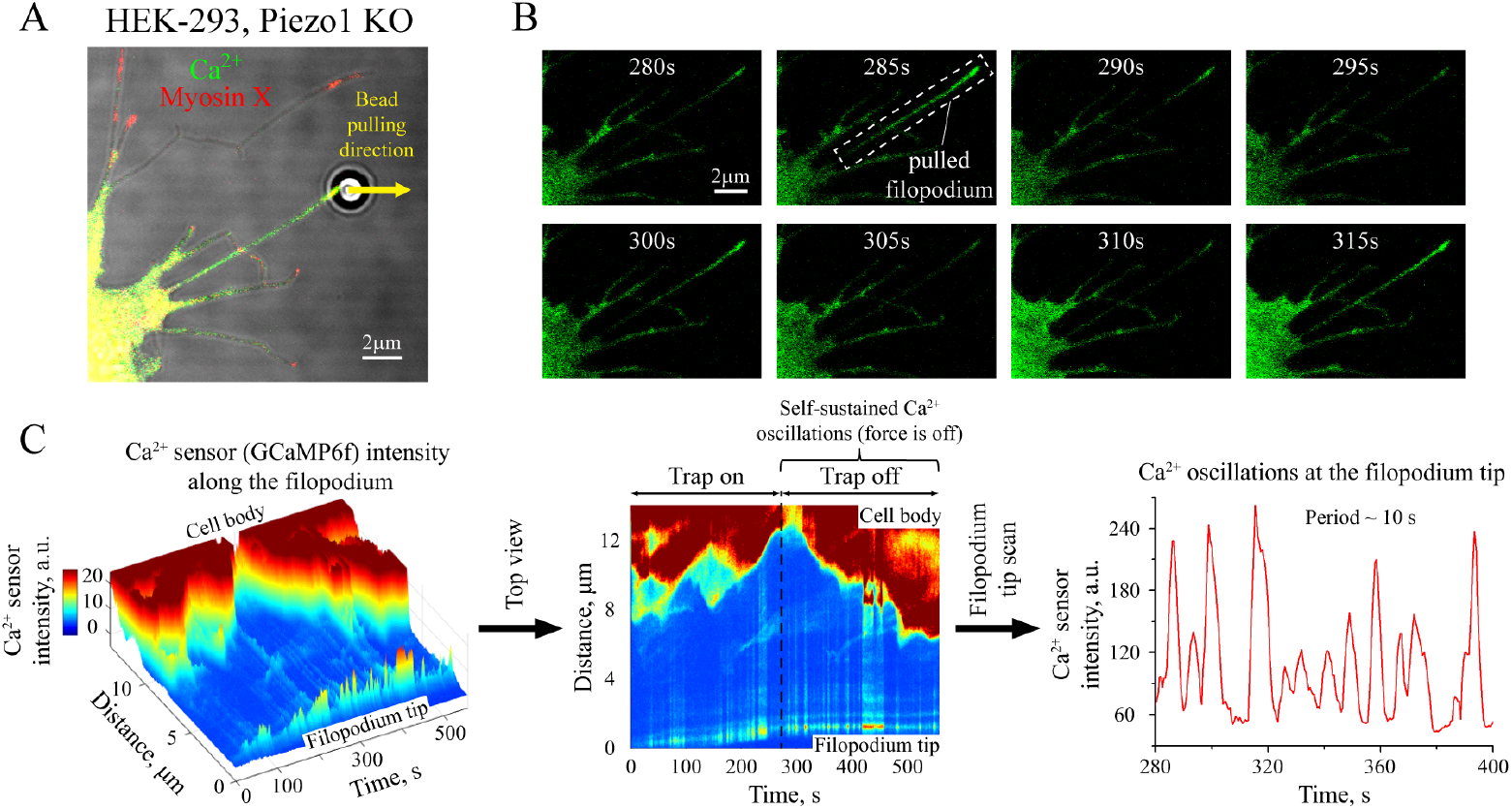
Quantification of the Ca^2+^ signaling in a mechanically stretched filopodium. **A.** A general view of a mechanically stretched filopodium. In the figure, mApple-myosin X is shown in red color; whereas, green color indicates Ca^2+^ sensor, GCaMP6f. **B.** Sequence of frames demonstrating change of the Ca^2+^ sensor intensity in the mechanically stretched filopodium, which is shown in panel A. **C.** Left 3D graph displays changes in the Ca^2+^ sensor intensity along the stretched filopodium as a function of time and distance from the filopodium tip. The middle panel shows a top view on the 3D graph from the left panel. Intra-filopodial Ca^2+^ oscillations in the form of periodic vertical strokes can be clearly seen in the graph. Once initiated, such Ca^2+^ oscillations keep going for several minutes even without further mechanical stretching of the filopodium when the optical trap is switched off. Right panel shows a representative time trace of the average Ca^2+^ sensor intensity at the filopodium tip, which reveals regular Ca^2+^ oscillations with a period of ~ 10 s.

Very similar force-dependent Ca^2+^ signals were also observed in MCF-7 and A2058 cells co-transfected with GCaMP6f and mApple-myosin X constructs, see Figure S2. Thus, we conclude that mechanical stretching of filopodia strongly promotes the appearance probability of filopodia-generated Ca^2+^ signals not only in HEK-293 cells, but also in other cell lines.

Force-dependent Ca^2+^ signals were also found to be produced by filopodia, whose growth was induced by expression of constitutively active Cdc42 (Q61L) in HEK-293 WT cells (see Movie 3), indicating that the observed phenomenon was rather general and not specific to myosin X-induced or Cdc42-induced filopodia.

We then checked whether such Ca^2+^ signals were induced by bona fide mechanosensitive calcium channels such as Piezo 1. Surprisingly, filopodial stretch-induced Ca^2+^ signals were also observed in a stable Piezo 1 knockout HEK-293 cell line (Figure 2). In addition, rescue of Piezo 1 via overexpression of Piezo 1-GFP construct in the knockout HEK-293 cells did not seem to have any effect on intra-filopodial Ca^2+^ signals as their behavior and time characteristics were similar to those found in HEK-293 Piezo 1 KO cells. Furthermore, HEK-293 cells are known to have a low expression level of Piezo 2 based on RNA-seq assays (39–41). Consistently, our qPCR analysis of HEK-293 Piezo 1 KO cell line demonstrated a very low level of Piezo 2 mRNA in comparison to housekeeping GAPDH gene, whose mRNA level was approximately 2^16^ ≈ 65000 times higher than that of Piezo 2 gene, see C_q_ values measured by qPCR for both of the genes in Table T1. Altogether, these results suggest that Piezo 1 and most probably Piezo 2 do not make a significant contribution to the observed force-dependent Ca^2+^ signals in filopodia.

Interestingly, intra-filopodial Ca^2+^ signals were also found to be independent from the integrin-mediated adhesion, since they were observed in filopodia stretched by using ConA-coated microbeads that do not activate integrin-related protein assembly at the attachment site (42, 43) (see Movie 4).

Finally, it should be noted that in ~ 40% (i.e., 4 out of N = 11) of experiments initial intra-filopodial Ca^2+^ signal resulted in strong elevation of the Ca^2+^ level in nearby cell cortex or even the whole cells during ~ 5 min observation period (see Movies 5A,B). On the other hand, the probability to see Ca^2+^ oscillations in the cell cytoplasm in the absence of filopodia stretching during a similar time interval was found to be < 10% (1 out of N = 11 cells). These observations suggest that the local mechanical activation of Ca^2+^ signaling in stretched filopodia may potentially trigger a global response in cells under certain conditions. Such phenomenon took place independently of the type of the beads (ConA- or fibronectin-covered) used in filopodia stretching experiments.

### Intra-filopodial Ca^2+^ signals are driven by Ca^2+^ influx

In order to understand whether the observed Ca^2+^ signals were caused by influx of Ca^2+^ ions from the cell culture medium or by Ca^2+^ release from endoplasmic reticulum, we treated cells with 1 μM thapsigargin for 1 hour, which caused release of Ca^2+^ ions from intracellular calcium-storage organelles. Should the Ca^2+^ signals in mechanically stretched filopodia originate from force-induced release of Ca^2+^ from intracellular calcium -storage organelles, such treatment would abolish intra-filopodial signaling as Ca^2+^ ions were already set free due to the action of the drug. In contrast to this expectation, we still observed elevation of the intra-filopodial Ca^2+^ level upon mechanical stretching of filopodia in thapsigargin-treated cells (~ 80%, 12 out of N = 15 cells) suggesting that the signal was likely caused by influx of Ca^2+^ ions from the exterior culture medium rather than from intracellular calcium-storage compartments, see Figure 3A-C and Movie 6.

**Figure 3.**
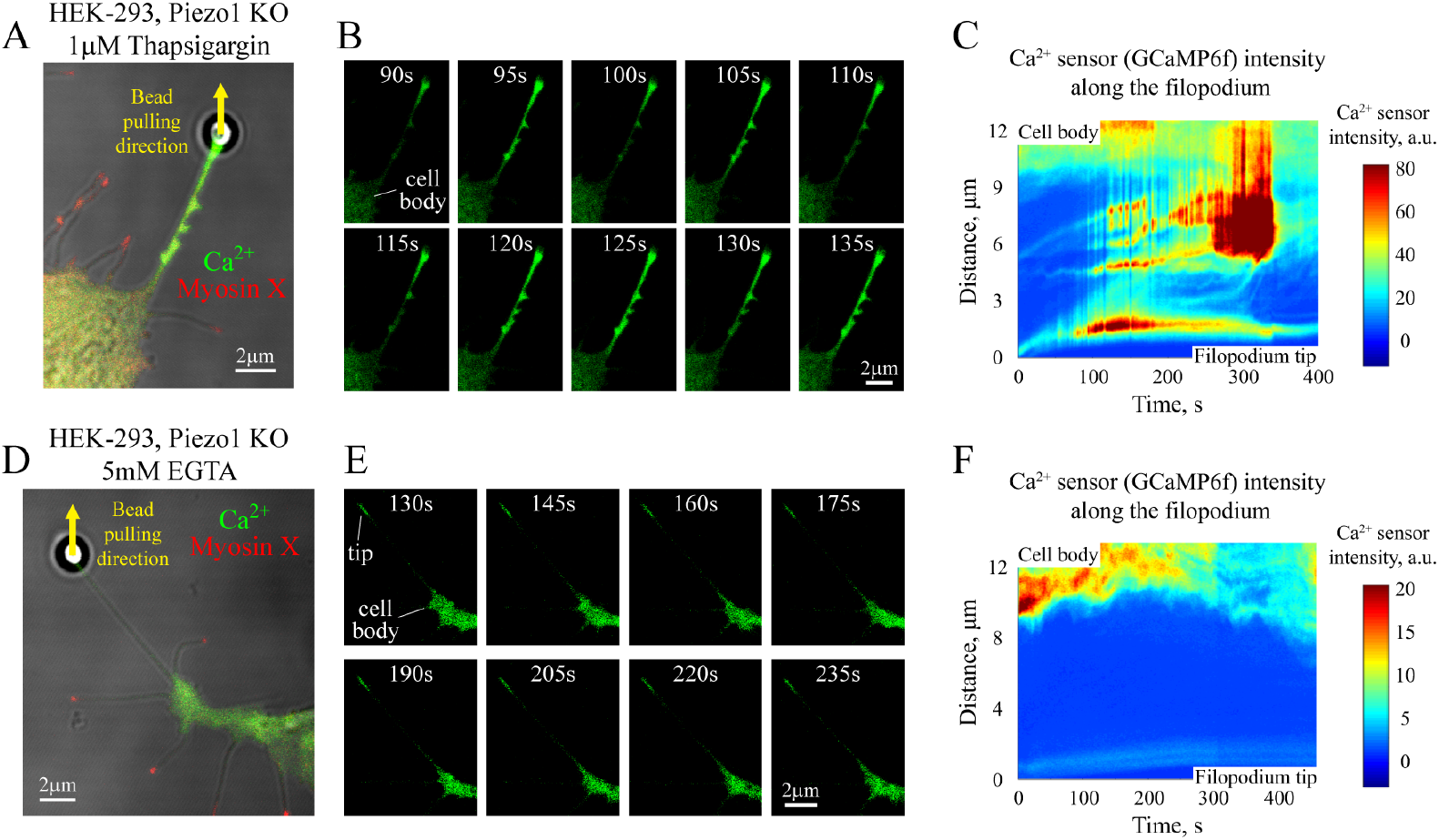
Intra-filopodial Ca^2+^ signals are driven by Ca^2+^ influx. **A.** View of a mechanically stretched filopodium of a HEK-293 Piezo 1 KO cell treated with 1 μM thapsigargin for 1 hr. **B.** Sequence of frames showing changes of the Ca^2+^ sensor (GCaMP6f) intensity in the mechanically stretched filopodium, which is displayed in panel A. **C**. Heatmap of the Ca^2+^ sensor intensity as a function of time and position on the filopodium, which is shown in panels A and B. Strong Ca^2+^ signal caused by filopodia stretching and its time-dependent oscillations can be clearly seen from the graph as well as from the frames shown in panel B. **D.** View of a mechanically stretched filopodium of a HEK-293 Piezo 1 KO cell in Ca^2+^-free cell culture medium in the presence of 5 mM EGTA. **E.** Sequence of frames demonstrating changes of the Ca^2+^ sensor (GCaMP6f) intensity in the mechanically stretched filopodium, which is shown in panel D. **F**. Heatmap of the intensity of Ca^2+^ sensor as a function of time and position on the filopodium, which is shown in panels D and E. As can be seen from the graph and frames presented in panel E, no Ca^2+^ signals have been observed inside the filopodium shaft upon mechanical stretching in the absence of free Ca^2+^ ions in the cell culture media. In panels A and D, mApple-myosin X is shown in red color, and Ca^2+^ sensor, GCaMP6f, is indicated in green color.

To test this hypothesis, we further performed filopodia stretching experiments in Ca^2+^-free cell culture medium, adding to it Ca^2+^ chelating agent, EGTA, at 5 mM concentration. Such experimental conditions led to complete disappearance of force-induced Ca^2+^ signals in the shafts of mechanically stretched filopodia of HEK-293 Piezo1 KO cells (in 10 out of N = 10 tested filopodia, 100%), see Figures 3D-F and Movie 7. These results suggest that intra-filopodial Ca^2+^ signals are indeed caused by influx of extracellular Ca^2+^ ions from the cell culture medium.

### L-type Ca^2+^ channels contribute to generation of intra-filopodial Ca^2+^ signals

Influx of Ca^2+^ ions into filopodia from the cell culture media indicated that some transmembrane Ca^2+^ channels were opened by filopodia mechanical stretching. The fact that intra-filopodial Ca^2+^ signals were observed in HEK-293 Piezo 1 KO cells, which have a very low expression level of Piezo 2, argues against possibility of Piezo 1 and Piezo 2 Ca^2+^ channels to be major candidates. In our effort to identify protein complexes responsible for such Ca^2+^ signals, we screened a few more previously reported mechanosensitive Ca^2+^ channels by using pharmacological inhibition assays in filopodia stretching experiments, checking whether there are any changes in the force-dependent Ca^2+^ signal response.

First, we tested TRPV4 mechanosensing Ca^2+^ channel (44, 45). To this aim, we treated HEK-293 Piezo 1 KO cells with 1 μM GSK2193874 for 30 min to inhibit TRPV4 (46). By using optical tweezers, it then has been found that despite the cells’ treatment with the drug, intra-filopodial Ca^2+^ signals still were present in 100% (12 out of N = 12) of mechanically stretched filopodia (see Movie 8). Thus, it seemed that TRPV4 channels did not substantially contribute to intra-filopodial Ca^2+^ signaling based on the cells treatment with the drug – a result, which is consistent with the recent finding showing that mammalian TRP channels, including TRPV4, cannot be activated by membrane stretching (47).

Next, we tested voltage-gated L-type Ca^2+^ channels since they have been previously shown to play an important role in formation of filopodia by promoting and stabilizing their growth (29). It has been previously reported that native L-type Ca^2+^ currents recorded in rat cardiomyocytes as well as human intestinal smooth muscle and rat mesenteric arterial smooth muscle cells demonstrate response to the cells’ stretching (48–50). Thus, the same channels may have been involved in the mechanically induced intra-filopodial Ca^2+^ influx observed in filopodia pulling experiments.

Indeed, qPCR assay performed on HEK-293 Piezo 1 KO cells has shown that the average mRNA level of CACNA1C gene, which encodes the pore-forming α1C subunit of L-type Ca^2+^ channels, is more than 10 times higher than that of Piezo 2 (see Table T1), suggesting that L-type Ca^2+^ channels may potentially have a stronger effect on the cell mechanosensing. Consistently, by treating HEK-293 Piezo 1 KO cells with 10 μM amlodipine besylate, a known inhibitor of L-type Ca^2+^ channels (51), for 30 min, we have found that the fraction of cells in which mechanically-induced Ca^2+^ signal was observed in stretched filopodia was significantly decreased. Namely, a strong Ca^2+^ signal was found to be produced only in approximately ~ 56% of tested filopodia (13 out of N = 23), which was lower than in the case of untreated HEK-293 Piezo 1 KO and HEK-293 WT cells, where the Ca^2+^ signal has been found to form in majority of mechanically stretched filopodia (~ 82%, 9 out N = 11, WT cells, and 100%, 11 out of N = 11, Piezo 1 KO cells). Figures 4A-F show two representative examples, one with a strongly suppressed Ca^2+^ signal in the presence of 10 μM amlodipine besylate in the cell culture medium (Figures 4A-C and Movie 9), and another with a Ca^2+^ signal retained under the same experimental conditions (Figures 4D-F and Movie 10).

**Figure 4.**
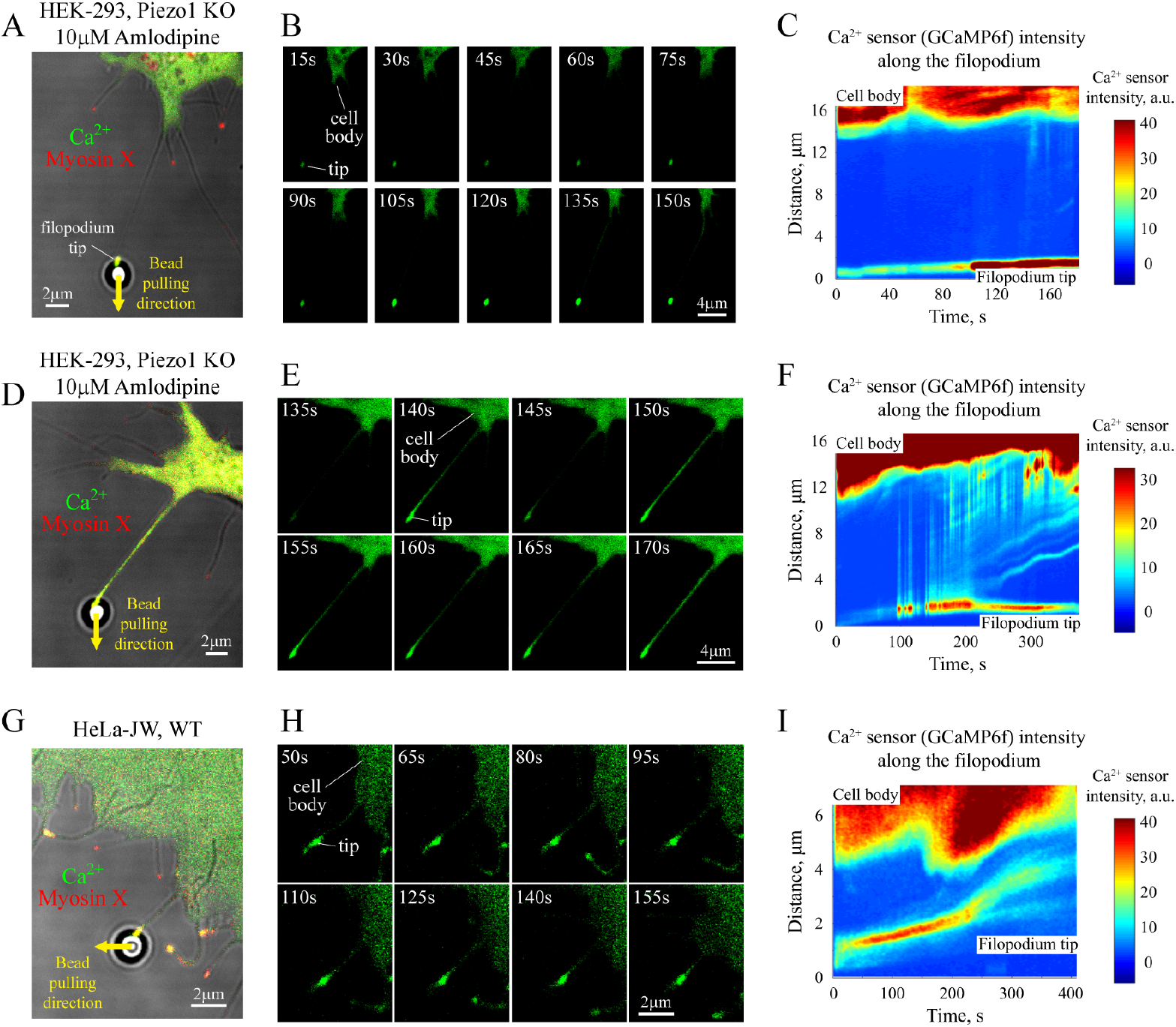
Intra-filopodial Ca^2+^ signaling depends on L-type Ca^2+^ channels. A, D. View of mechanically stretched filopodia of two HEK-293 Piezo 1 KO cells in the presence of 10 μM amlodipine besylate in the cell culture medium – one with Ca^2+^ signal being suppressed by the drug (panel A), and another with Ca^2+^ signal being retained despite the presence of the drug (panel D). **G.** View of a mechanically stretched filopodium of a HeLa-JW WT cell. **B, E, H.** Sequence of frames demonstrating changes in the intensity of Ca^2+^ indicator, GCaMP6f, in the mechanically stretched filopodia shown in panels A, D and G, respectively. **C, F, I.** Heatmaps of the Ca^2+^ sensor intensity as a function of time and position on the filopodium that correspond to the cells presented in panels A, D and G, respectively. In panels A, D and G, mApple-myosin X is shown in red color, and Ca^2+^ sensor, GCaMP6f, is indicated in green color.

To check whether suppression of the Ca^2+^ signal by amlodipine besylate was reversible, we washed out the drug from the medium and repeated filopodia stretching experiments after 30 minutes incubation time. It has been found that the percentage of filopodia generating Ca^2+^ signal upon mechanical stretching (100%, 13 out of N = 13 tested filopodia) returned to a level similar to that observed in untreated cells (~ 90%, 9 out of N = 10 tested filopodia). These results strongly suggest that suppression of Ca^2+^ signal in stretched filopodia was indeed reversible and caused by amlodipine besylate.

While the exact reason why amlodipine besylate affected Ca^2+^ influx only in a fraction of the treated cells was unclear, overall the above data indicate that L-type Ca^2+^ channels are a key player being involved in formation and propagation of Ca^2+^ signals in mechanically stretched filopodia.

To further test this hypothesis, we carried out additional filopodia stretching experiments on HeLa cells that have been previously reported to have a low expression level of L-type Ca^2+^ channels (29, 40, 41). Consistently, the force-induced Ca^2+^ influx was detected in less than half of the pulled filopodia of HeLa-JW cells, with only ~ 42% of filopodia (8 out N = 19) generating sufficiently strong Ca^2+^ signal, see Figures 4G-I and Movie 11. This result was in stark contrast to ~ 80-100% probability to observe Ca^2+^ signal in mechanically stretched filopodia of HEK-293 cells. After transfection of HeLa-JW cells with a plasmid encoding the pore-forming α1C subunit (CaV1.2) of L-type Ca^2+^ channels, the probability to spot the force-induced Ca^2+^ signal in stretched filopodia increased significantly – up to ~ 79% (15 out of N = 19 of tested filopodia, see also Movie 12). On the other hand, almost no change in the Ca^2+^ signal appearance probability (~ 36%, 5 out of N = 14 tested filopodia) has been found in mechanically stretched filopodia of HeLa-JW cells transfected with an empty vector backbone. Altogether, these results strongly suggest that L-type Ca^2+^ channels play a critical role in formation of the force-induced Ca^2+^ signals in filopodia.

### Probability of Ca^2+^ influx into mechanically stretched filopodia under different experimental conditions

In order to have a more quantitative and statistically unbiased understanding of how the observed Ca^2+^ signal in mechanically stretched filopodia is affected by various factors and to summarize the results presented above, we measured the maximum intensity of Ca^2+^ signal in the shafts of stretched filopodia (excluding the tip region attached to the trapped bead), which was then normalized to the average Ca^2+^ sensor intensity in the cell cytoplasm, see Methods and Figure S3 for more details. The results obtained from different cell types under various experimental conditions are displayed in the form of a boxplot in Figure 5.

**Figure 5.**
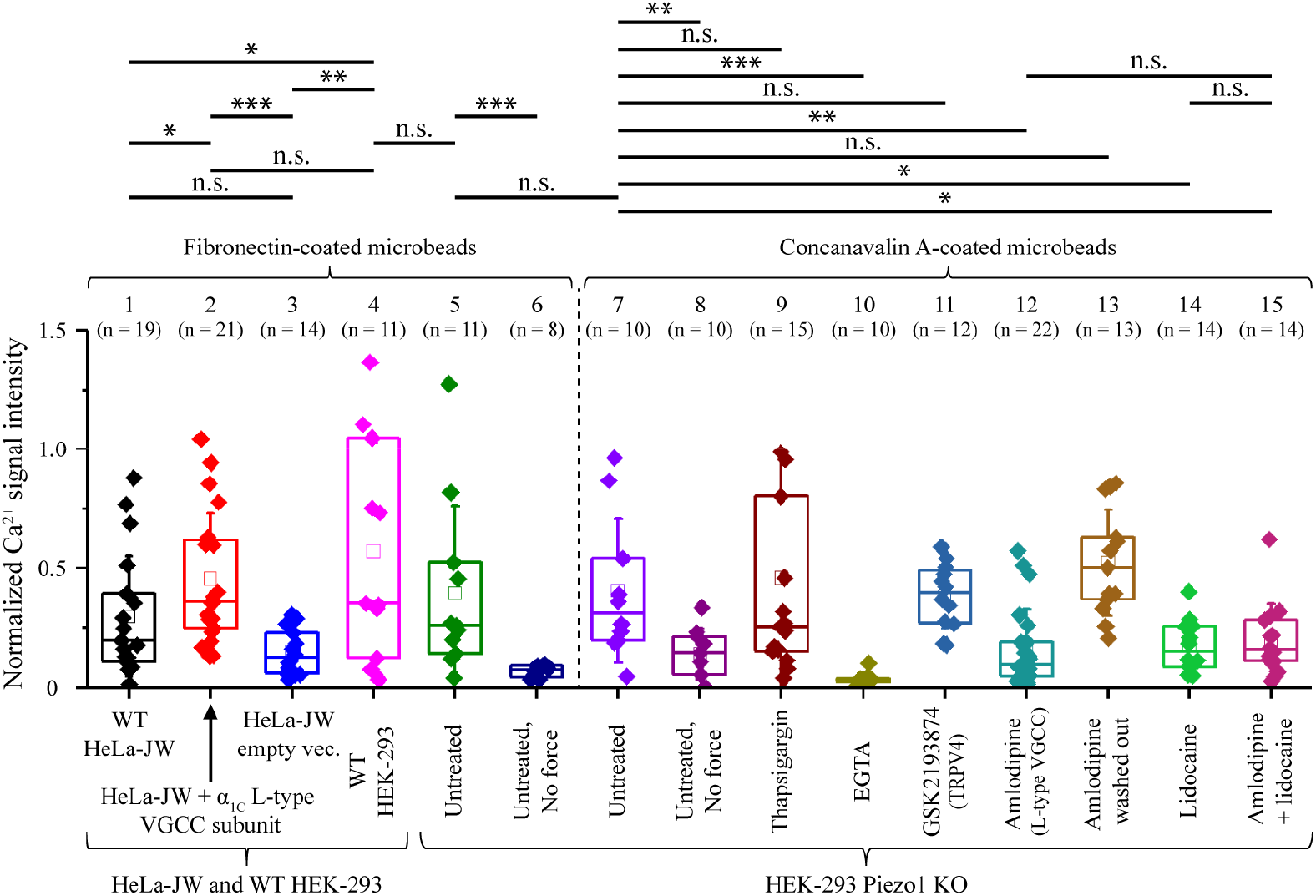
Maximum normalized intensities of Ca^2+^ signals measured in mechanically stretched filopodia under different experimental conditions. Data shown in the figure indicate that mechanical stretching of filopodia as well as presence of EGTA or L-type Ca^2+^ channels’ inhibitor (amlodipine besylate) in the cell culture media – all have very significant impact on the intra-filopodial Ca^2+^ signaling. The bars over the boxplot display pairwise statistical difference between the columns (n.s. – non-significant, * – p < 0.05, ** – p < 0.01, *** – p < 0.001). The p-values were calculated by using non-parametric Mann-Whitney test. For each indicated experimental condition, the data were obtained based on N = 8-22 filopodia stretching experiments (see numbers in parentheses shown above the columns) performed on 8-15 different cells – i.e., in the majority of the cases filopodia were taken from different cells. Unless otherwise indicated in the graph captions, maximum strength of the Ca^2+^ signal was measured in the presence of stretching force applied to tested filopodia. The original data are provided in Data S1.

As can be seen from the figure, Ca^2+^ signals produced by bead-attached filopodia in the absence of mechanical load had very low normalized maximum intensities (~ 0.1–0.2, see columns 6 and 8) corresponding to the peak levels of individual weak transient Ca^2+^ spikes. In contrast, in mechanically stretched filopodia (columns 4, 5 and 7), Ca^2+^ influx was much stronger and occurred at higher frequency independently of the expression level of Piezo 1 Ca^2+^ channels in HEK-293 cells and the microbeads’ coating used in experiments, resulting in significantly larger normalized maximum intensity of the Ca^2+^ signal. Use of Ca^2+^-free cell culture medium with additional 5 mM EGTA completely abolished all intra-filopodial Ca^2+^ signals (column 10). In contrast, no statistically significant change in the maximum normalized Ca^2+^ sensor intensity has been observed in mechanically stretched filopodia of HEK-293 Piezo 1 KO cells treated with 1 μM of thapsigargin (column 9) in comparison to untreated cells (column 7). Thus, it can be concluded that the force-induced Ca^2+^ signals in mechanically stretched filopodia emerge due to influx of extracellular Ca^2+^ rather than due to Ca^2+^ release from calcium-storing cell organelles, such as endoplasmic reticulum.

Data obtained in the presence of 1 μM of TRPV4 channel inhibitor, GSK2193874, reveal that this mechanosensitive Ca^2+^ channel does not make a substantial contribution to the observed intra-filopodial Ca^2+^ signals (column 11). In contrast, cells treatment with 10 μM amlodipine besylate, a known inhibitor of L-type voltage-gated Ca^2+^ channels (VGCC), as well as subsequent wash out of the drug from the cell culture medium had significant effects on the maximum normalized intensities of Ca^2+^ signals that form in response to filopodia stretching (columns 12 and 13). These results indicate that L-type Ca^2+^ channels make a significant contribution to the observed force-dependent Ca^2+^ signaling in filopodia.

Indeed, by carrying out filopodia stretching experiments on wild-type (WT) HeLa cells (column 1), which have been reported to have a low expression level of L-type Ca^2+^ channels based on RNA-seq assay (29, 40, 41), it has been found that the maximum normalized intensities of force-induced Ca^2+^ signals produced by filopodia of such cells are weaker than in the case of WT HEK-293 cells (column 4) whose expression level of L-type Ca^2+^ channels is higher based on the same RNA-seq assay (40, 41). Furthermore, after transfection of HeLa-JW cells with a plasmid encoding the pore-forming α1C subunit (CaV1.2) of L-type Ca^2+^ channels, the maximum normalized intensities of Ca^2+^ signals generated by mechanically stretched filopodia increased significantly (column 2), reaching practically the same level as in the case of WT HEK-293 cells (column 4). As such increase has not been observed in control experiments done on HeLa-JW cells transfected with an empty vector backbone (column 3), it can be concluded that L-type Ca^2+^ channels indeed play a critical role in formation of the observed force-dependent Ca^2+^ signals in filopodia.

Finally, it is interesting to note that treatment of HEK-293 Piezo 1 KO cells with 2 mM lidocaine, a known inhibitor of voltage-gated Na^+^ channels (52), or with a combination of 2 mM lidocaine and 10 μM amlodipine besylate resulted in decrease of the maximum normalized intensities of intra-filopodial Ca^2+^ signals (columns 14 and 15 in Figure 5), which was similar to the case of cells treated with amlodipine besylate alone (column 12). These experimental data suggest existence of an intricate interplay between the membrane potential and voltage-gated L-type Ca^2+^ channels in the force-dependent generation of Ca^2+^ influx into mechanically stretched filopodia.

### Force-dependent Ca^2+^ influx triggers local calpain activity

Previous experimental studies done on neurite growth cones suggest that Ca^2+^ signals generated by filopodia are used by neural cells during the growth cone pathfinding process, which appears to be driven by local Ca^2+^-dependent activation of calpain protease (30, 53). The latter is known to be involved in regulation of many adhesion-related proteins, resulting in local changes of the cell adhesion strength to extracellular matrix (ECM), which is one of the molecular mechanisms underlying cells motion through ECM (34). Based on these studies and our experimental findings it can be hypothesized that the force-dependent Ca^2+^ influx may potentially affect calpain activity in mechanically stretched filopodia, leading to downstream regulation of cell adhesion complexes.

To test whether the observed intra-filopodial Ca^2+^ influx is sufficient to activate calpain, we performed filopodia stretching experiments on HEK-293 Piezo1 KO cells transfected with a vector encoding CFP-YFP FRET sensor for calpain activity. This sensor consists of CFP and YFP fluorescent domains linked by a peptide containing a calpain cleavage site (54). Upon activation, calpain cleaves the peptide linking CFP and YFP domains, resulting in loss of the apparent FRET signal, which can be easily detected in experiments (i.e., the ratio between YFP and CFP fluorescence intensities decreases).

Figures 6A-B demonstrate typical time-dependent changes in the apparent FRET ratio of the CFP-YFP sensor in two different cells in response to mechanical stretching of the filopodia, which are indicated by cyan arrows. From these figures it can be seen that calpain activation usually takes place within the first 1-2 minutes after the start of filopodia stretching, which is consistent with a local elevation of the Ca^2+^ concentration in filopodia due to the force-induced Ca^2+^ influx. Interestingly, Figure 6B shows that application of mechanical force not only results in activation of calpain in the stretched filopodium itself, but also frequently causes increase in the calpain activity in nearby cell regions. This phenomenon is likely related to propagation of the initial intra-filopodial Ca^2+^ signal induced by the filopodium stretching to nearby lamellipodia or even the whole cell body that has been mentioned at the end of the first Result section.

**Figure 6.**
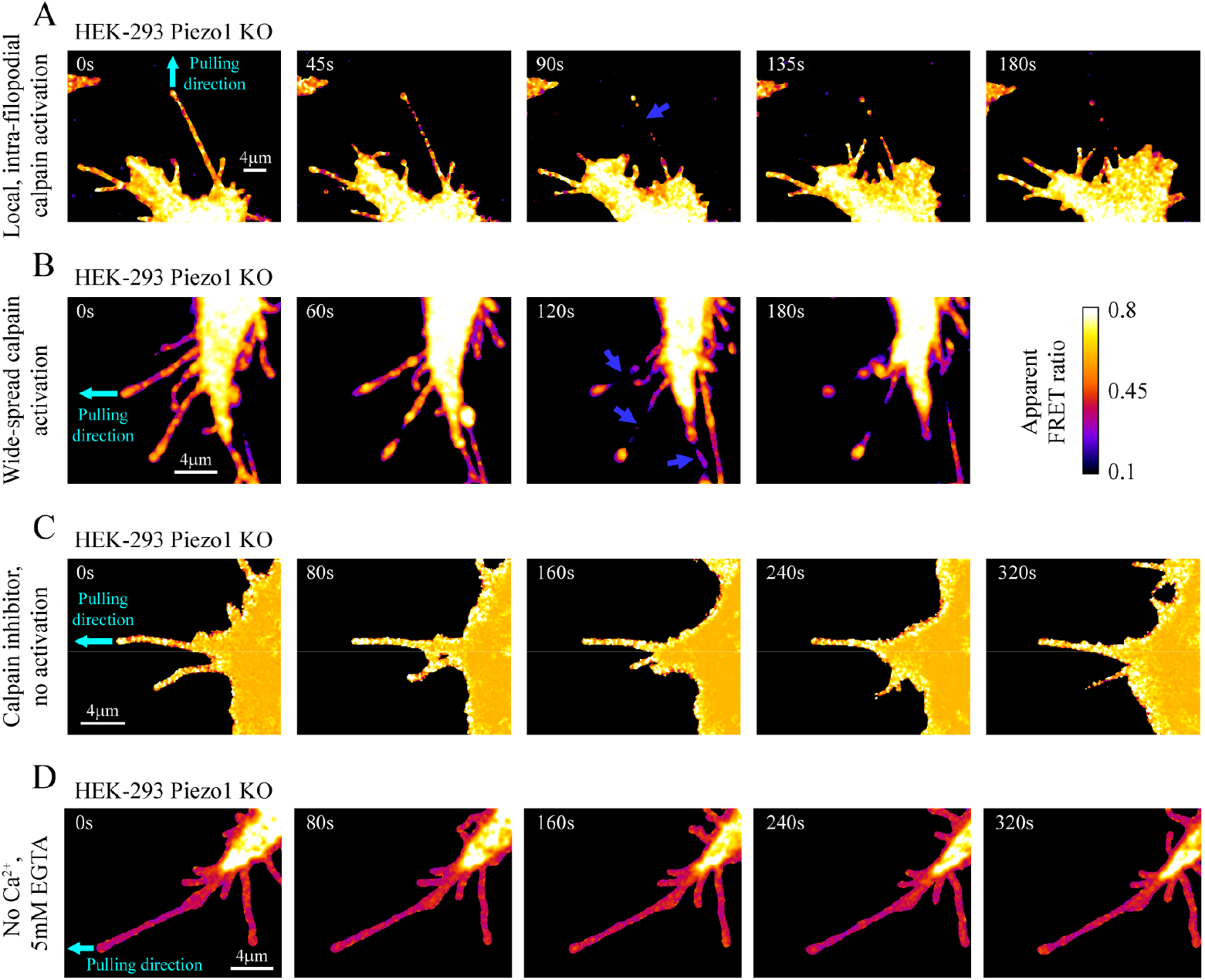
Force-dependent Ca^2+^ influx promotes calpain activation in living cells. **A.** Sequence of frames showing local decrease in the apparent FRET ratio of CFP-YFP calpain activity sensor in a stretched filopodium. Starting from T = 90 sec, local disappearance of the FRET signal (marked by the dark blue arrow) can be observed in the stretched filopodium. **B.** Sequence of frames demonstrating global activation of calpain in response to mechanical stretching of a filopodium. Disappearance of the FRET signal in the stretched filopodium as well as in nearby filopodia indicated by the dark blue arrows can be clearly seen at T = 120 sec. **C.** Sequence of frames showing the effect of 100 μM calpain inhibitor, ALLN. It can be seen from the figure that ALLN completely abolishes force-induced activation of calpain in the stretched filopodium, which is indicated by the stable FRET signal coming from the CFP-YFP calpain activity sensor. **D.** Sequence of frames demonstrating time-dependent FRET signal coming from calpain activity sensor in a mechanically stretched filopodium in Ca^2+^-free cell culture medium. Stability of the FRET signal suggests that calpain is not activated in response to the filopodium stretching in the absence of free Ca^2+^ in solution. In panels A-D, cyan arrows indicate stretched filopodia and their pulling directions. In all four panels, filopodia stretching was commenced at T = 0 sec.

To check that the observed decrease in the FRET signal coming from the CFP-YFP sensor is caused by calpain activation, we treated cells with 100 μM calpain inhibitor, ALLN (55), for 1 hour and repeated the above experiment. It has been found that the FRET signal in mechanically stretched filopodia of such cells remained constant over minutes of the force application for all the tested filopodia (N = 6) (see Figure 6C), confirming that the observed FRET signal decrease in the case of untreated cells is indeed caused by calpain activation in stretched filopodia.

To find out whether calpain activation takes place due to influx of extracellular Ca^2+^, we repeated filopodia stretching experiments in Ca^2+^-free cell culture media, which additionally contained 5 mM EGTA. Under such experimental conditions, FRET signal coming from the CFP-YFP calpain activity sensor stayed at the same stable level without showing any tendency to decrease over ~ 5-6 minutes of observation period for all the stretched filopodia (N = 6), see Figure 6D. In combination with previous results, this finding suggests that Ca^2+^ influx into filopodia is necessary for calpain activation. Thus, calpain activation in response to filopodia stretching is one of the main downstream effects of the force-induced intra-filopodial Ca^2+^ influx, which has a strong impact on local adhesion strength of living cells (34).

## Discussion

In this study, it has been shown that filopodia are highly sensitive structures producing local Ca^2+^ signals in response to mechanical load. Application of pulling forces in the range of tens of pN to filopodia tips was found to dramatically increase influx of extracellular Ca^2+^ through transmembrane channels, resulting in formation of persistent intra-filopodial Ca^2+^ oscillations. The latter could last for many minutes even after the force was released, indicating existence of filopodia memory effect. Such a force-dependent activation of Ca^2+^ signaling in filopodia has been observed in several distinct types of living cells, including human embryonic kidney cells (HEK-293), breast cancer epithelial cells (MCF-7) and metastatic melanoma cells (A2058), suggesting that it is based on rather universal molecular mechanisms. However, in some other cell types (HeLa-JW) force-dependent Ca^2+^ influx into filopodia was found to be weak if existing at all.

To obtain insights into the potential role of various Ca^2+^ channels in this phenomenon, we utilized pharmacological inhibition and knock-out assays based on HEK-293 cell line in combination with filopodia stretching experiments. In this way, we excluded the possibility that such force-dependent activation of Ca^2+^ influx into filopodia is mediated by Piezo 1 or TRPV4 transmembrane proteins, which are two known mechanosensitive Ca^2+^ channels (31, 32). Furthermore, qPCR assay performed on HEK-293 Piezo 1 KO cells showed that Piezo 2 mechanosensing Ca^2+^ channel is also unlikely to be an important component in the observed phenomenon as its transcription level was found to be very low.

Besides the above mechanosensitive Ca^2+^ channels, GPR68, a G protein-coupled receptor, is also known to induce elevation of intracellular Ca^2+^ level in living cells upon mechanical stimulation (56). However, GPR68-related mechanism is based on the release of Ca^2+^ ions from intracellular storages sensitive to thapsigargin treatment (56), which cannot be the cause of the force-dependent extracellular Ca^2+^ influx into filopodia observed in our study.

On the other hand, results of pharmacological inhibition of L-type Ca^2+^ channels in HEK-293 cells demonstrated that these channels play a major role in the force-induced Ca^2+^ signaling in stretched filopodia. This finding is further supported by observation that overexpression of the pore-forming α1C subunit of L-type Ca^2+^ channels in HeLa-JW cells that normally express it weakly, leads to a statistically significant increase in the Ca^2+^ influx in response to filopodia stretching.

However, it is not yet clear whether these channels function as primary mechanosensors or amplify an upstream signal generated by other type of molecules. In this connection, it is interesting to note that treatment of HEK-293 Piezo 1 KO cells with lidocaine, which inhibits membrane depolarization by Na^+^ channels (52), leads to a similar level of decrease in strength of intra-filopodial Ca^2+^ signals as inhibition of L-type Ca^2+^ channels by amlodipine besylate. As it is known that some Na^+^ channels are mechanosensitive (57), there remains a possibility that primary mechano-response is generated by mechanosensing Na^+^ channels, while voltage-gated L-type Ca^2+^ channels function downstream of them. Other potentially mechanosensory molecules, such as adhesion- and actin cytoskeleton-related talin and formin, which are found in filopodia (7, 58–64), could in principle also be primary initiators of mechano-response. Of note, however, integrin-dependent mechanosensitivity does not seem to be necessary for mechano-stimulation of Ca^2+^ influx into filopodia in our system. Indeed, Ca^2+^ influx could be induced by applying force not only to fibronectin-coated microbeads that specifically interact with integrins, but also to microbeads coated with ConA, which interact with any molecules bearing α-D-mannosyl and α-D-glucosyl groups and which traditionally used as a negative control in integrin-signaling studies (42, 43).

Thus, our findings suggest existence of either direct or indirect mechanosensitivity of L-type Ca^2+^ channels residing on the filopodia surface. Prior to this work, the potential mechanosensitivity of L-type Ca^2+^ channels was only implicated in muscle cells, such as rat cardiomyocytes (50), human intestinal smooth muscle (48) and rat mesenteric arterial smooth muscle cells (49), mainly from studies based on whole-cell patch-clamp experiments. However, our study has shown that these Ca^2+^ channels may contribute to mechanosensing behavior of a much larger group of living cells and revealed importance of filopodia in this type of mechano-response. In addition, while previous studies showed that L-type Ca^2+^ channels localized to filopodia are important for maintenance of filopodia integrity (29), here we demonstrated that these channels as well mediate the force-induced Ca^2+^ influx into filopodia.

Furthermore, we have shown that force-dependent Ca^2+^ influx into mechanically stretched filopodia through transmembrane channels is sufficient for activation of Ca^2+^-sensitive calpain protease. By performing filopodia stretching experiments on cells expressing calpain activity sensor, we found that application of pulling force results in strong activation of calpain protease inside the stretched filopodia within a short time period of 1-2 minutes. Moreover, such calpain activation was completely abolished after removal of free Ca^2+^ from the cell culture medium, suggesting that the force-induced Ca^2+^ influx through filopodial transmembrane Ca^2+^ channels is absolutely necessary for it. These results are in good agreement with previous experimental studies showing that elevation of the intracellular Ca^2+^ level leads to a rapid activation of calpain protease within a short time period of 30–60 s, which results in degradation of the calpain target proteins within the next 1–2 minutes (65, 66).

Interestingly, previous studies show that μ- and m-calpain sensitivity to Ca^2+^ is strongly enhanced in the presence of PIP2 phospholipids *in vivo* (65, 67). It is also known that filopodia formation is typically initiated on PIP2 lipid rafts, which are required for binding of several essential filopodial proteins such as IRSp53 and Ena/VASP (2). Thus, filopodia are likely to be enriched with PIP2 phospholipids, which would explain high sensitivity of calpain protease to the force-induced Ca^2+^ influx into stretched filopodia observed in our study.

What could be possible biological functions of the force-induced Ca^2+^ influx into filopodia? One simple consideration is based on known targets of calpain proteolytic activity. Calpain cleaves talin (68) and therefore can in principle release the integrin talin-mediated adhesion of filopodia tips. Thus, if a filopodium is pulled via integrin-mediated contact, activation of calpain by Ca^2+^ could disrupt such a contact and release the tension that may potentially harm the filopodium and/or cell, consistently with the role of L-type Ca^2+^ channels in maintenance of filopodia integrity (29). In this connection, it is also interesting that an “exceptional” cell type, HeLa-JW, in which filopodia stretching induced very weak, if any, Ca^2+^ entry, demonstrates another response to mechanical tension – force-induced elongation of filopodia (7) that can also prevent filopodia rupture.

Calpain, however, can produce more complex effects on integrin adhesions rather than simply disrupt them. In particular, in growth cone filopodia, calpain-mediated cleavage of talin and FAK results in inhibition of both adhesions’ formation and their disassembly. This affects axon growth characteristics, such as repulsive turning and response to the substrate rigidity (69). Moreover, besides calpain, Ca^2+^ influx can regulate other local targets in filopodia, including calcineurin phosphatase, which is involved in a variety of signaling pathways (70). Thus, mechanically induced Ca^2+^ influx can trigger different signaling cascades, via calpain and other targets, which could affect growth and adhesion of filopodia. In addition, we have found that in some cases Ca^2+^ signal can spread from filopodia to the cell body. Such signal propagation and its possible function deserves further investigation.

In summary, our study clearly demonstrates that Ca^2+^ influx into filopodia is a basic signaling response to physiological level of tensile forces applied to filopodia tips and therefore can be used by cells in different processes of mechano-orientation and motion guidance, involving response to the matrix rigidity and topography. Moreover, we established that in cells of different types such response is mediated by L-type Ca^2+^ channels rather than known mechanosensing Ca^2+^ channels like TRPV4, and Piezo 1 and Piezo 2. Involvement of L-type Ca^2+^ channels in different type of mechanosensory mechanisms is an interesting avenue for the future studies.

## Materials and Methods

### Cell lines, plasmids and inhibitors

Wild-type HEK-293T, MCF-7 and A2058 cells used in this study were obtained from ATCC company. HEK-293T Piezo1 KO stable cell line was kindly provided by the Boris Martinac’s and Ardem Patapoutian’s laboratories. HeLa-JW, a subline of a HeLa cervical carcinoma cell line derived in the laboratory of J. Willams (Carnegie-Mellon University, USA) on the basis of better attachment to plastic dishes (71), was obtained from the laboratory of B. Geiger (72).

The calcium sensor (pGP-CMV-GCaMP6f) used in this study was a gift from Douglas Kim & GENIE Project (Addgene plasmid # 40755; http://n2t.net/addgene:40755; RRID:Addgene_40755). pCMV-calpainsensor plasmid was a gift from Isabelle Richard (Addgene plasmid # 36182; http://n2t.net/addgene:36182; RRID:Addgene_36182). The plasmid containing the α1C subunit (CaV1.2) of L-type Ca^2+^ channels was a gift from Diane Lipscombe (Addgene plasmid # 26572; http://n2t.net/addgene:26572; RRID:Addgene_26572). The mApple-myosin X was subcloned by the Protein Cloning and Expression Core facility of the MBI.

All cell lines were cultured in DMEM media supplemented with sodium pyruvate and 10% Hi-FBS (Life Technologies). To express protein constructs in living cells, jetPrime transfection reagent was used to introduce plasmids into the cells a day before the experiment. Four hours later after the transfection, cells were re-plated onto 4-well glass bottom dishes coated with fibronectin at a concentration of 10 μg/ml. After that cells were incubated overnight in DMEM media, allowing them to firmly attach to the surface of the fibronectin-coated dishes. On the day of the experiment, the cell culture medium was switched to 1X Ringer’s balanced salt buffer supplemented with 11mM glucose. pH level of all the cell culture media used in experiments was in 7.4–7.6 range.

For sarco/endoplasmic reticulum Ca^2+^-ATPase (SERCA) inhibition studies, cells were treated with 1 μM thapsigargin (abcam, ab120286) for 1 hr before the experiment. For TRPV4 Ca^2+^ channel inhibition studies, 1 μM GSK2193874 (Sigma-Aldrich, SML0942) was added to cells 30 min prior the experiment. Inhibition of voltage-gated L-type Ca^2+^ channels was done by introduction of 10 μM amlodipine besylate (Tocris, catalog # 2571) into the cell culture medium 30 min before the experiment. For inhibition of voltage-gated Na^+^ channels, cells were pre-treated with 2 mM lidocaine (Sigma-Aldrich, L7757) for 30 min prior the start of the experiment. Finally, inhibition of calpain protease was achieved by adding 100 μM ALLN (Peptide International, IAL– 3671-PI) to the cell culture medium before the experiment. In all experiments, inhibitors remained in the medium during the entire observation period.

### Optical tweezers experiments

Detailed description of the optical tweezers setup and experimental procedures can be found in ref. (7). Calibration of the optical trap stiffness, *k,* was done by using the viscous flow calibration method (73). The measured value of *k* was 0.51 ± 0.07 pN/nm. The force, *F*, applied to stretched filopodia in optical tweezers experiments was obtained by using the formula: *F* = *k*Δ*x*, where *k* is the stiffness of the optical trap, and Δ*x* is the measured deviation of the bead center from the axis of the optical trap (see Figure 1C).

### RT-qPCR experiments

For quantification of the transcription level of Piezo 2, CACNA1C and GAPDH genes, total RNA of cultured HEK-293 Piezo 1 KO cells were extracted by using Qiagen RNeasy Micro Kit (Cat No./ID: 74004). Then 5 μg of the extracted RNA was reversely transcribed by utilizing Tetro cDNA Synthesis Kit (Bioline, BIO-65043) and random Hexamer Primer according to the manufacturer’s protocols. The RT-qPCR reactions were carried out on a Biorad C1000 thermo cycler by using custom-synthesized Taqman Probes for Piezo 2 (Hs00926218_m1), CACNA1C (Hs00167681_m1) and GAPDH (Hs99999905_m1) genes, and Taqman Gene Expression Mastermix from Thermofisher. Obtained C_q_ values of the RT-qPCR runs were then used for analysis.

### Data processing

To quantify intensities of the filopodia fluorescent signals coming from Ca^2+^ sensor, GCaMP6f, the “plot profile” tool of ImageJ program was used in the study. For this purpose, in each movie the axis of a mechanically stretched filopodium was selected as a contour along which intensity of the signal was measured. To prevent distortion of the signal by the background noise, the average noise level (which was typically < 0.3 a.u.) was subtracted from each frame of the collected experimental movies. Furthermore, to ensure consistency of the measurements the following steps were taken during filopodia stretching experiments. First, we studied only those filopodia, which were lying on the horizontal surface of a glass coverslip. As filopodia are rather thin membrane protrusions (typical diameter < 500nm), this makes it possible to image a whole filopodium in a single Z stack. Next, during experiments, Z-axis drift of the focal plane of the microscope was minimized by monitoring and compensating the deviation of the imaging plane from the position corresponding to the sharpest filopodia contrast in the bright field (see, for example, Figure S3A). This allowed us to eliminate even small drifts along Z-axis of the microscope (~ 200 nm) as the filopodia bright field image has strong sensitivity on the change in Z-position of the microscope focal plane.

Experimental measurements indicate that by using such a compensation mechanism, all of the changes in the intensity of Ca^2+^ sensor in the filopodium shaft due to the focal plane drift along Z-axis direction can be reduced to a very low level of ≤ 1.5-2 a.u. over an experimentally relevant timescale of several minutes. As in all of the experiments with mechanically stretched filopodia the average signal intensity upon Ca^2+^ influx activation was typically > 10 a.u., it can be concluded that the relative error in intensity measurements due to the microscope focal plane movement was ≤ 15 %. This makes it possible to accurately quantify the role of extracellular forces in activation of Ca^2+^ influx into mechanically stretched filopodia.

As for X and/or Y microscope stage movements that were necessary to generate a pulling force on filopodia, to minimize potential measurement errors that may arise from the horizontal movements, we selected a region of interest (ROI) of 50 pixels around filopodia contour by setting the line width = 50 pixels in the “plot profile” tool of ImageJ, and ensured that filopodia remained within the monitored ROI during the whole process of stretching, see Figure S3B. This method makes it possible to accurately quantify intensity profiles of filopodia independently of the microscope stage movements in both X and Y directions.

Finally, in order to eliminate potential bias in data processing, all of the filopodial Ca^2+^ sensor signals were normalized by the average intensity of a cell cytoplasmic region of 10-20 μm^2^ size that was measured prior to filopodia stimulation by force, see Figure S3B.

## Data availability

All study data are included in this article and SI Appendix.

## Acknowledgements

We would like to thank Dr. Naila Omar Khayyam Alieva (IMCB, A*STAR, Singapore), Dr. Kate Poole (UNSW, Australia) and Dr. Johanna Ivaska (University of Turku, Finland) for encouraging and productive discussions. We also thank Dr. F. Margadant, Peng Qiwen, Liu Jun, Ong Hui Ting, Bi Feng and Mak Kah Jun (MBI Microscopy Core facility), and Dr. Virgile Viasnoff for their kind help with the optical tweezers. The research was funded by the National Research Foundation (NRF), Prime Minister’s Office, Singapore under its NRF Investigatorship Programme (NRF Investigatorship Award No. NRF-NRFI2016-03, to J.Y.), the National Research Foundation, Prime Minister’s Office, Singapore and the Ministry of Education under the Research Centres of Excellence programme (to J.Y., M.P.S., A.D.B.), Singapore Ministry of Education Academic Research Fund Tier 3 (MOE2016-T3-1-002 to J.Y., M.P.S., A.D.B., in part), and by the Singapore Ministry of Education Academic Research Fund Tier 2 (MOE2018-T2-2-138 to A.D.B.).

## Author contributions

A.K.E. and M.Y. designed the study and performed the experiments; A.K.E. analyzed the data; A.K.E., M.Y. and J.Y. interpreted the data. A.K.E., M.Y. and J.Y. wrote the paper; M.P.S., A.D.B. and B.M. provided cell lines and plasmids as well as helpful insights into the studied molecular systems; A.K.E. and J.Y. supervised the research.

## Supplementary figures

**Figure S1.**
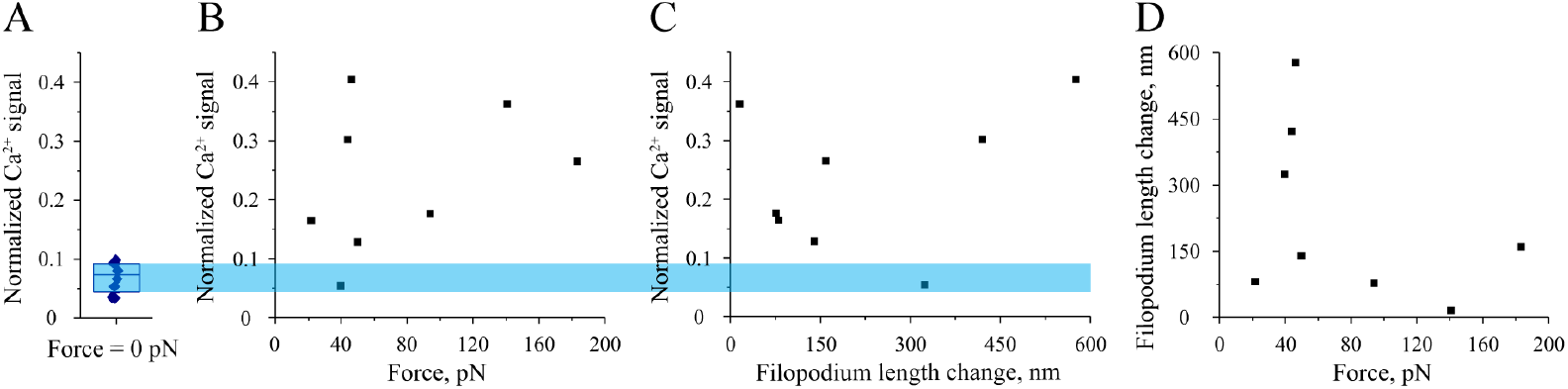
**A.** Normalized maximum intensity of the Ca^2+^ sensor measured in filopodia attached to fibronectin-covered microbeads in the absence of mechanical load (F = 0 pN). **B.** Normalized maximum intensity of the Ca^2+^ sensor vs mechanical load measured in filopodia stretching experiments performed on WT HEK-293 cells. **C.** Normalized maximum intensity of the Ca^2+^ sensor vs force-induced filopodia extensions (i.e., strain) measured in the same experiments as in panel A. **D.** Filopodia extensions plotted vs applied mechanical load. Data points shown in the graphs A-D were collected at the moment of appearance of the first force-induced Ca^2+^ signal in pulled filopodia. In panels A-C, intensity of the Ca^2+^ sensor fluorescent signal, which was observed in the filopodia shafts (excluding filopodia tip regions), is normalized to the intensity of the Ca^2+^ sensor in the cell cytoplasm measured prior to filopodia stretching (if any), see more details in Method section. Blue shaded area in panels AC shows the range of Ca^2+^ signal intensities corresponding to panel A, which were measured in the absence of mechanical load.

**Figure S2.**
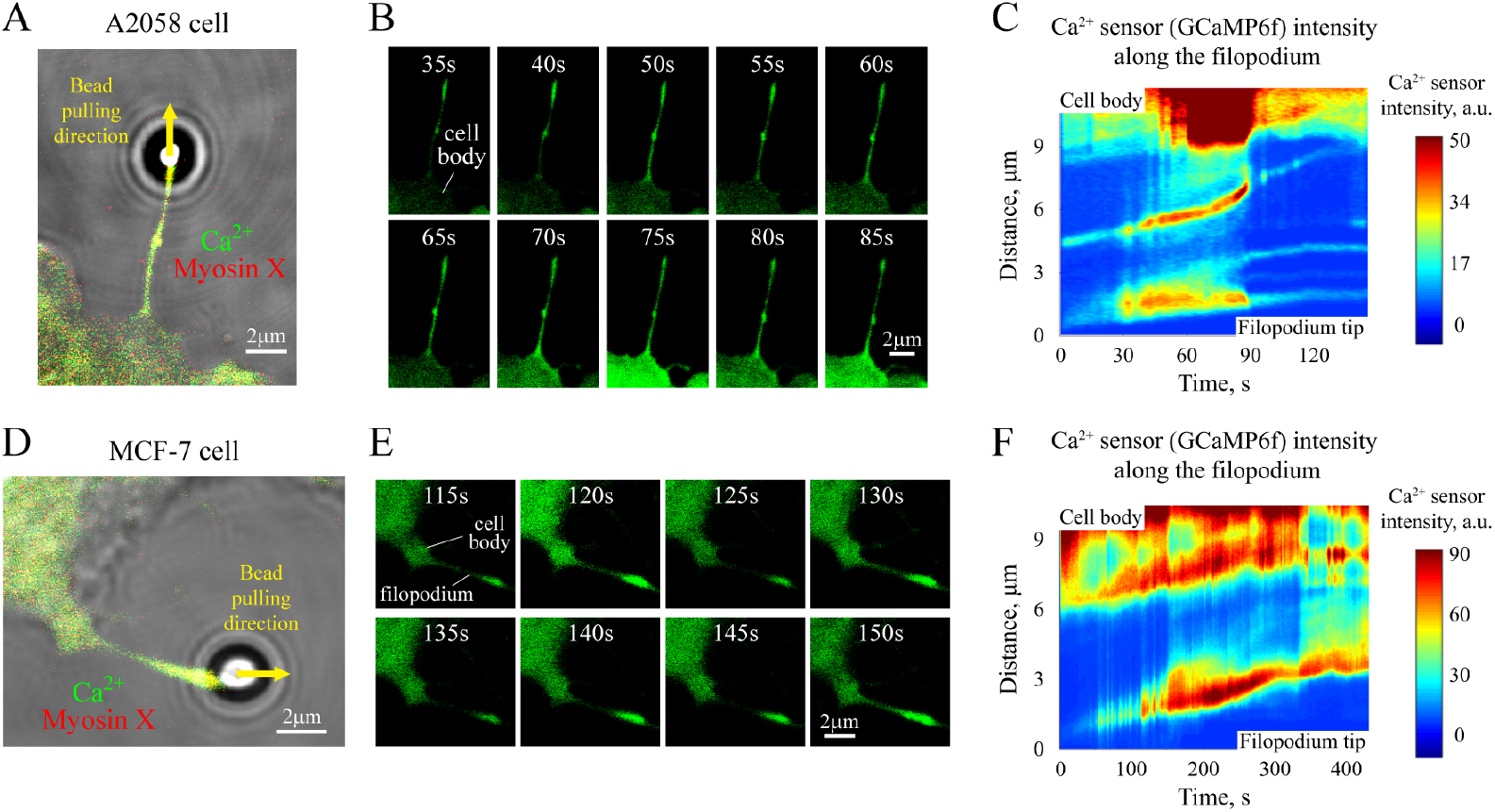
Ca^2+^ signaling in a mechanically stretched filopodia of A2058 and MCF-7 cells. A, D. View of mechanically stretched filopodia of A2058 and MCF-7 cells. **B, E.** Sequence of frames demonstrating changes in the intensity of Ca^2+^ indicator, GCaMP6f, in the mechanically stretched filopodia shown in panels A and D, respectively. **C, F.** Heatmaps of the Ca^2+^ sensor intensity as a function of time and position on the filopodium that correspond to the cells presented in panels A and D, respectively. In panels A and D, mApple-myosin X is shown in red color, and Ca^2+^ sensor, GCaMP6f, is indicated in green color.

**Figure S3.**
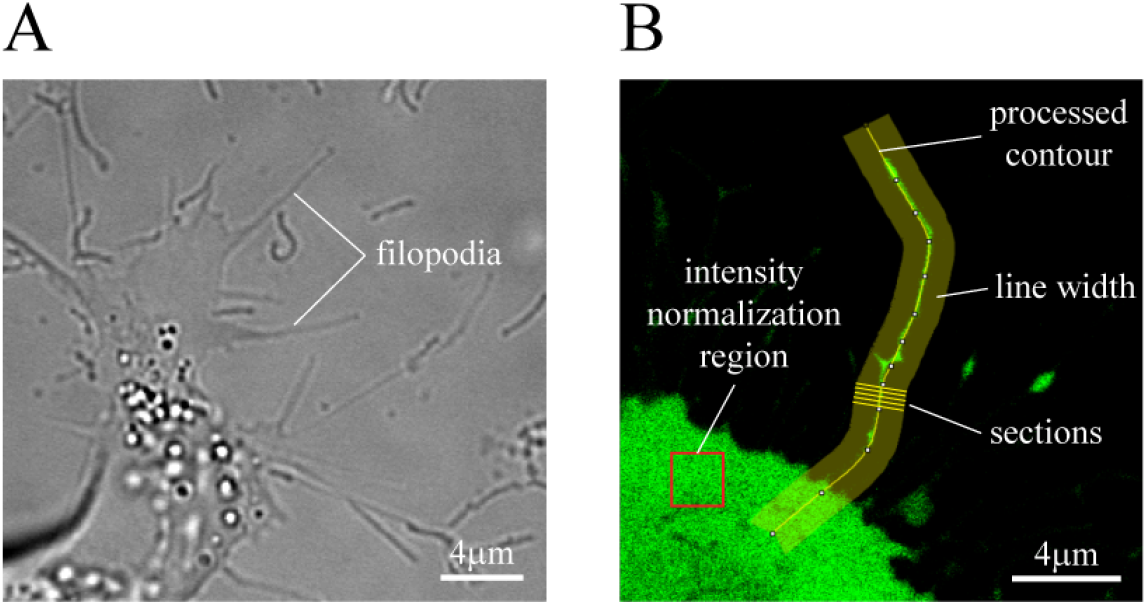
**A.** Position of the microscope focal plane corresponding to the sharpest images of filopodia in the bright field. **B.** Data processing procedure. To obtain an intensity profile of a stretched filopodium, the “plot profile” tool of ImageJ was used. It calculates the average intensity of sections of a fixed width (50 pixel) that are perpendicular to the contour of a stretched filopodium (shown by yellow curve). Several such sections are schematically shown in panel B. To represent data in an unbiased way, all intensity profiles of stretched filopodia were normalized to the average intensity of a cell cytoplasmic region of ~ 10-20 μm^2^ size (red rectangle in panel B) located outside of the region occupied by the cell nucleus.

**Table T1.** C_q_ values indicating the relative mRNA levels of GAPDH housekeeping gene (control), Piezo 2 gene, and CACNA1C gene (encodes the pore-forming α_1C_ subunit of L-type Ca^2+^ channels) measured in qPCR assay, which was done for three different samples prepared from HEK-293 Piezo 1 KO cells.

| Gene | C<sub>q</sub>, sample 1 | C<sub>q</sub>, sample 2 | C<sub>q</sub>, sample 3 |
| --- | --- | --- | --- |
| GAPDH (control) | 20.43 | 20.21 | 20.46 |
| Piezo 2 | 37.11 | 35.72 | 36.10 |
| CACNA1C | 32.41 | 32.18 | 32.09 |

## Supplementary movies

### Movie 1

The movie shows a fibronectin-coated microbead attached to the surface of a force-unloaded (i.e., optical trap is off) myosin X-induced filopodium of a HEK-293 cell. The bead demonstrates persistent movement along the filopodium towards the cell body, with no Ca^2+^ signal being observed inside the bead-attached filopodium. The left panel in the movie displays composite frames that combine the bright field, green (GCaMP6f Ca^2+^ sensor) and red (mApple-myosin X) channels; whereas, the right panel demonstrates only the green channel (GCaMP6f Ca^2+^ sensor). The duration of the original video clip is 4 min 22 s. The movie was recorded at 2 fps rate and displayed at 75 fps.

### Movies 2A,B

The movies show a mechanically stretched myosin X-induced filopodium of a wild-type HEK-293 cell. The force was applied to the tip region of the filopodium by using an optically trapped fibronectin-coated microbead (see Figure 1C for details). Strong Ca^2+^ signal produced by the stretched filopodium in response to the applied mechanical load can be clearly seen in both movies. The left panel in Movie 2A displays composite frames that combine the bright field, green (GCaMP6f Ca^2+^ sensor) and red (mApple-myosin X) channels; whereas, the right panel demonstrates only the green channel (GCaMP6f Ca^2+^ sensor). Original duration of Movie 2A is 5 min 30 s. The movie was recorded at 2 fps rate and displayed at 75 fps. As for Movie 2B, it demonstrates a set of frames from Movie 2A corresponding to the moment of the Ca^2+^ signal appearance in the stretched filopodium in slow motion (displayed at 5 fps rate).

### Movie 3

The movie shows a mechanically stretched filopodium induced by constitutively active GFP-Cdc42 (Q61L) in a wild-type HEK-293 cell. The force was applied to the tip of the filopodium by using an optically trapped fibronectin-coated microbead. Clearly visible Ca^2+^ signal inside the stretched filopodium can be seen in the movie above the background level generated by GFP-Cdc42. The left panel in the movie displays composite frames that combine the bright field and the green channel (GCaMP6f Ca^2+^ sensor + GFP-Cdc42); whereas, the right panel demonstrates only the green channel (GCaMP6f Ca^2+^ sensor + GFP-Cdc42). The duration of the original video clip is 7 min 16 s. The movie was recorded at 2 fps rate and displayed at 120 fps.

### Movie 4

The movie shows a mechanically stretched myosin X-induced filopodium of a HEK-293 Piezo 1 KO cell. The force was applied to the tip region of the filopodium by using an optically trapped concanavalin A-coated microbead. Strong oscillating Ca^2+^ signal can be clearly seen in the movie inside the stretched filopodium. The left panel in the movie displays composite frames that combine the bright field, green (GCaMP6f Ca^2+^ sensor) and red (mApple-myosin X) channels; whereas, the right panel demonstrates only the green channel (GCaMP6f Ca^2+^ sensor). The duration of the original video clip is 3 min 28 s. The movie was recorded at 2 fps rate and displayed at 60 fps.

### Movies 5A,B

The movies show a mechanically stretched myosin X-induced filopodium of a HEK-293 Piezo 1 KO cell. The force was applied to the tip of the filopodium by using an optically trapped concanavalin A-coated microbead. Activation of the cell cortex at the base of the stretched filopodium caused by increase in the intra-filopodial Ca^2+^ level can be clearly seen in Movie 5A at T = 472 s. The left panel in Movie 5A displays composite frames that combine the bright field, green (GCaMP6f Ca^2+^ sensor) and red (mApple-myosin X) channels; whereas, the right panel demonstrates only the green channel (GCaMP6f Ca^2+^ sensor). The duration of the original video clip is 10 min 4 s. The movie was recorded at 2 fps rate and displayed at 120 fps. As for Movie 5B, it demonstrates a set of frames from Movie 5A corresponding to the moment of the cell cortex activation in slow motion (displayed at 10 fps rate).

### Movie 6

The movie shows a mechanically stretched myosin X-induced filopodium of a HEK-293 Piezo 1 KO cell in the presence of 1 μM thapsigargin (sarco/endoplasmic reticulum Ca^2+^-ATPase inhibitor) in the cell culture medium. The force was applied to the tip of the filopodium by using an optically trapped concanavalin A-coated microbead. Strong force-induced Ca^2+^ signal can be clearly seen inside the stretched filopodium. The left panel in the movie displays composite frames that combine the bright field, green (GCaMP6f Ca^2+^ sensor) and red (mApple-myosin X) channels; whereas, the right panel demonstrates only the green channel (GCaMP6f Ca^2+^ sensor). The duration of the original video clip is 5 min 27 s. The movie was recorded at 2 fps rate and displayed at 90 fps.

### Movie 7

The movie shows a mechanically stretched myosin X-induced filopodium of a HEK-293 Piezo 1 KO cell in Ca^2+^-free cell culture medium in the presence of 5 mM EGTA. The force was applied to the tip of the filopodium by using an optically trapped concanavalin A-coated microbead. As can be seen from the movie, presence of Ca^2+^-chelating EGTA in the medium completely abolishes Ca^2+^ signals in the shaft of the mechanically stretched filopodium. The left panel in the movie displays composite frames that combine the bright field, green (GCaMP6f Ca^2+^ sensor) and red (mApple-myosin X) channels; whereas, the right panel demonstrates only the green channel (GCaMP6f Ca^2+^ sensor). The duration of the original video clip is 4 min 42 s. The movie was recorded at 2 fps rate and displayed at 75 fps.

### Movie 8

The movie shows a mechanically stretched myosin X-induced filopodium of a HEK-293 Piezo 1 KO cell in the presence of 1 μM GSK2193874 (TRPV4 Ca^2+^ channel inhibitor) in the cell culture medium. The force was applied to the tip of the filopodium by using an optically trapped concanavalin A-coated microbead. Strong intra-filopodial Ca^2+^ signal resulting in the global cell activation can be clearly seen in the movie. The left panel in the movie displays composite frames that combine the bright field, green (GCaMP6f Ca^2+^ sensor) and red (mApple-myosin X) channels; whereas, the right panel demonstrates only the green channel (GCaMP6f Ca^2+^ sensor). The duration of the original video clip is 2 min 0 s. The movie was recorded at 2 fps rate and displayed at 30 fps.

### Movie 9

The movie shows a mechanically stretched myosin X-induced filopodium of a HEK-293 Piezo 1 KO cell in the presence of 10 μM amlodipine besylate (L-type Ca^2+^ channels’ inhibitor) in the cell culture medium. The force was applied to the tip of the filopodium by using an optically trapped concanavalin A-coated microbead. As can be seen from the movie, inhibition of L-type Ca^2+^ channels results in suppression of the force-induced Ca^2+^ signal in the stretched filopodium. This type of behavior was observed in ~ 44% of the studied cells. The left panel in the movie displays composite frames that combine the bright field, green (GCaMP6f Ca^2+^ sensor) and red (mApple-myosin X) channels; whereas, the right panel demonstrates only the green channel (GCaMP6f Ca^2+^ sensor). The duration of the original video clip is 3 min 4 s. The movie was recorded at 2 fps rate and displayed at 45 fps.

### Movie 10

The movie shows a mechanically stretched myosin X-induced filopodium of a HEK-293 Piezo 1 KO cell in the presence of 10 μM amlodipine besylate (L-type Ca^2+^ channels’ inhibitor) in the cell culture medium. The force was applied to the tip of the filopodium by using an optically trapped concanavalin A-coated microbead. This video demonstrates an example of an amlodipine-treated cell which retained Ca^2+^ signaling in the mechanically stretched filopodium despite the presence of the L-type Ca^2+^ channels’ inhibitor in the cell culture medium. Such type of behavior was observed in ~ 56% of the treated cells. The left panel in the movie displays composite frames that combine the bright field, green (GCaMP6f Ca^2+^ sensor) and red (mApple-myosin X) channels; whereas, the right panel demonstrates only the green channel (GCaMP6f Ca^2+^ sensor). The duration of the original video clip is 5 min 48 s. The movie was recorded at 2 fps rate and displayed at 90 fps.

### Movie 11

The movie shows a mechanically stretched myosin X-induced filopodium of a wild-type HeLa-JW cell. The force was applied to the tip of the filopodium by using an optically trapped fibronectin-coated microbead. It can be seen from the movie that mechanical stretching of the filopodium results in generation of a very weak intra-filopodial Ca^2+^ signal. The left panel in the movie displays composite frames that combine the bright field and the green channel (GCaMP6f Ca^2+^ sensor); whereas, the right panel demonstrates only the green channel (GCaMP6f Ca^2+^ sensor). The duration of the original video clip is 3 min 16 s. The movie was recorded at 2 fps rate and displayed at 45 fps.

### Movie 12

The movie shows a mechanically stretched myosin X-induced filopodium of a HeLa-JW cell transfected with a plasmid encoding the pore-forming α_1c_ subunit (Cav1.2) of L-type Ca^2+^ channels. The force was applied to the tip of the filopodium by using an optically trapped fibronectin-coated microbead. Strong force-induced Ca^2+^ signal can be clearly seen inside the stretched filopodium in the movie. The left panel in the movie displays composite frames that combine the bright field and the green channel (GCaMP6f Ca^2+^ sensor); whereas, the right panel demonstrates only the green channel (GCaMP6f Ca^2+^ sensor). The duration of the original video clip is 2 min 58 s. The movie was recorded at 2 fps rate and displayed at 45 fps.

